# Systematic optimization and benchmarking of synchro-PASEF for high-throughput phosphoproteome profiling

**DOI:** 10.64898/2026.06.26.734570

**Authors:** Dain R. Brademan, Angelina A. Mullarkey, Mia L. Greeson, Sarah Szvetecz, Olga Vitek, Emily E. Blythe, Ruth Hüttenhain

## Abstract

High-throughput data-independent acquisition (DIA) workflows paired with short chromatographic separations are increasingly adopted for systems biology and clinical proteomics. However, narrower peak widths from rapid separations demand faster mass spectrometer cycle times to maintain quantitative depth and reproducibility. The synchro-PASEF acquisition mode on timsTOF mass spectrometers diagonally scans across ion mobility and *m/z* space, enabling efficient sampling of the precursor ion cloud with shortened cycle times. While synchro-PASEF has demonstrated competitive identification depth for global protein abundance samples compared to conventional dia-PASEF, its performance for phosphoproteomics–where the precursor ion cloud is characteristically broader and bimodally distributed–has not been evaluated. Here, we systematically optimized synchro-PASEF methods for phosphoproteomics and benchmarked performance against two dia-PASEF methods across three sub-hour separations.

We found that synchro-PASEF performance depends critically on balancing diagonal window number, total isolation width, and gradient length, with longer gradients favoring more windows for selectivity and shorter gradients favoring fewer windows to preserve sampling frequency. An optimized configuration quantified over 19,000 localized phosphosites using a 23-minute separation. Retention time summation (RTsum) with a factor of 2 increased phosphopeptide identifications by 5–20% and reduced phosphosite-level coefficients of variation by up to 30% across all dia-PASEF and synchro-PASEF methods tested. Using β2-adrenergic receptor (B2AR) activation as a signaling model, we demonstrate that label-free DIA phosphoproteomics can be used to model phosphoproteomics dose-response relationships, showing that synchro-PASEF and dia-PASEF produce highly concordant phosphoproteomic responses, with comparable numbers of responding phosphosites, similar effect sizes, and nearly identical predicted protein kinase A (PKA) substrates downstream of the activated B2AR. While synchro-PASEF did not surpass optimized dia-PASEF in identification depth, its comparable biological performance and amenability to post-acquisition optimization through RTsum support its utility for high-throughput phosphoproteomics. This work provides a transferable framework for synchro-PASEF method optimization and demonstrates the broad utility of retention time summation for PASEF-based phosphoproteomics workflows.

**Highlights:**

- Systematic benchmarking of synchro-PASEF for typical phosphoproteomics workflows.
- RT summation improves IDs and quantitative precision for dia-PASEF and synchro-PASEF
- dia-PASEF and synchro-PASEF capture dose-response phosphosignaling with comparable performance
- Provides transferable framework for high-throughput DIA method design

**Graphical Abstract:** 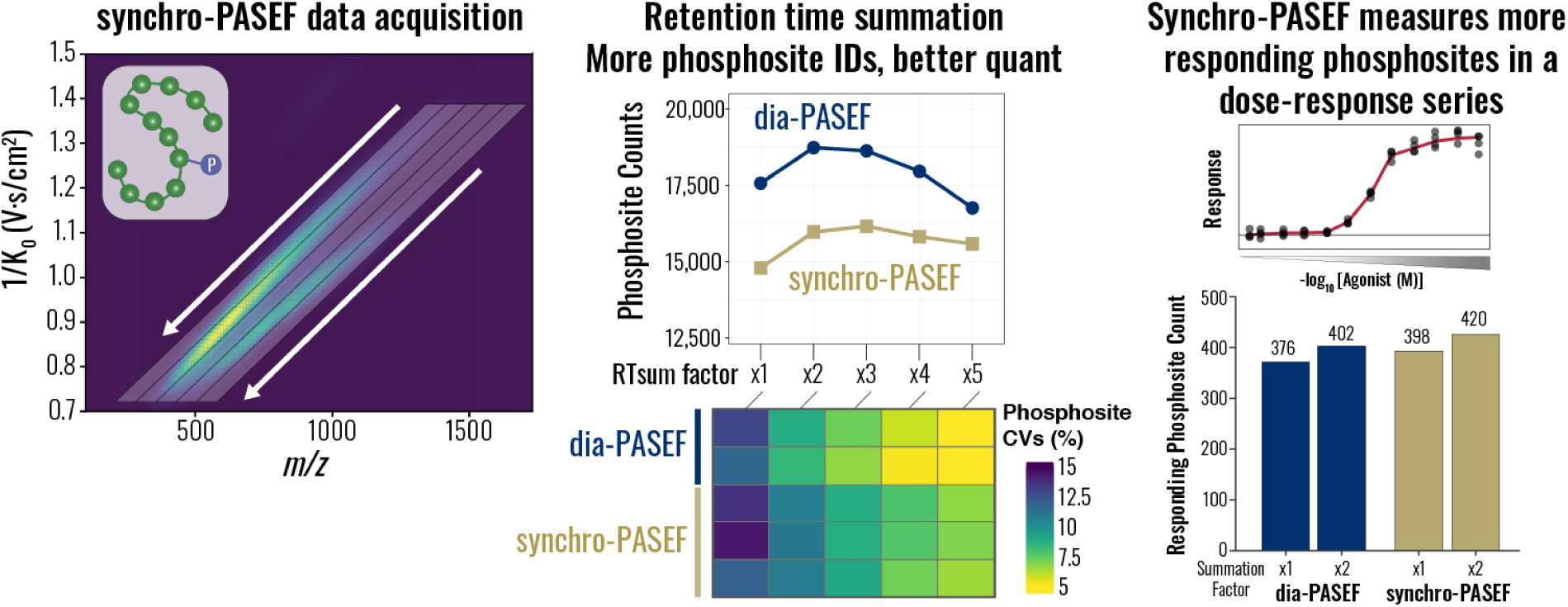

## Introduction

High-throughput mass spectrometry (MS)-based proteomics has become a cornerstone of systems biology and clinical research where large sample cohorts demand analytical strategies that balance depth, quantitative robustness, and acquisition speed. To meet these requirements, contemporary proteomic workflows frequently combine shortened chromatographic gradients with optimized MS acquisition methods, enabling comprehensive analysis of many samples with low instrumental overhead. (1–5) While shortened separations substantially increase throughput, they reduce chromatographic peak widths and the number of data points acquired across each elution profile, which can compromise quantitative precision and reproducibility–particularly for low-abundance species. (6–8) These limitations can, in principle, be mitigated by modifying the MS data acquisition strategy to increase sampling frequency of eluting peptide features. (9, 10) Common strategies include shortening the precursor accumulation time, acquiring spectra at a lower resolution to reduce transient lengths, or specifically in data-independent acquisition (DIA) workflows, using fewer and wider isolation windows for precursor fragmentation. (11–13)

Recent advances in trapped ion mobility spectrometry (TIMS) coupled with parallel accumulation serial fragmentation (PASEF) have enabled new acquisition modes designed to improve peptide ion utilization and sampling efficiency. (14) TIMS provides an additional dimension of separation of peptide ions based on gas phase collisional cross-section. The combination of DIA, TIMS, and PASEF–collectively referred to as dia-PASEF–has emerged as a widely adopted standard for high-throughput DIA proteomics on timsTOF instruments. (15, 16) In dia-PASEF acquisition, peptide ions are accumulated and separated in the TIMS device by collisional cross section, then sequentially released in mobility-resolved packets. During each TIMS ramp, the analytical quadrupole steps through defined precursor *m/z* windows, enabling data-independent fragmentation across the diagonally correlated precursor ion mobility-*m/z* area in a stair-like manner (**Fig. 1A)**. A limitation of this stepped quadrupole isolation scheme samples only a subset of the precursor ion population per TIMS ramp due to set quadrupole isolation, and ion utilization efficiency is reduced where DIA isolation windows do not track the diagonal precursor ion cloud well in sparse regions of the mobilogram. (17, 18) Computational optimization of window placement and width can partially address this limitation by better matching isolation boundaries to the precursor ion density. (17)

**Figure 1:**
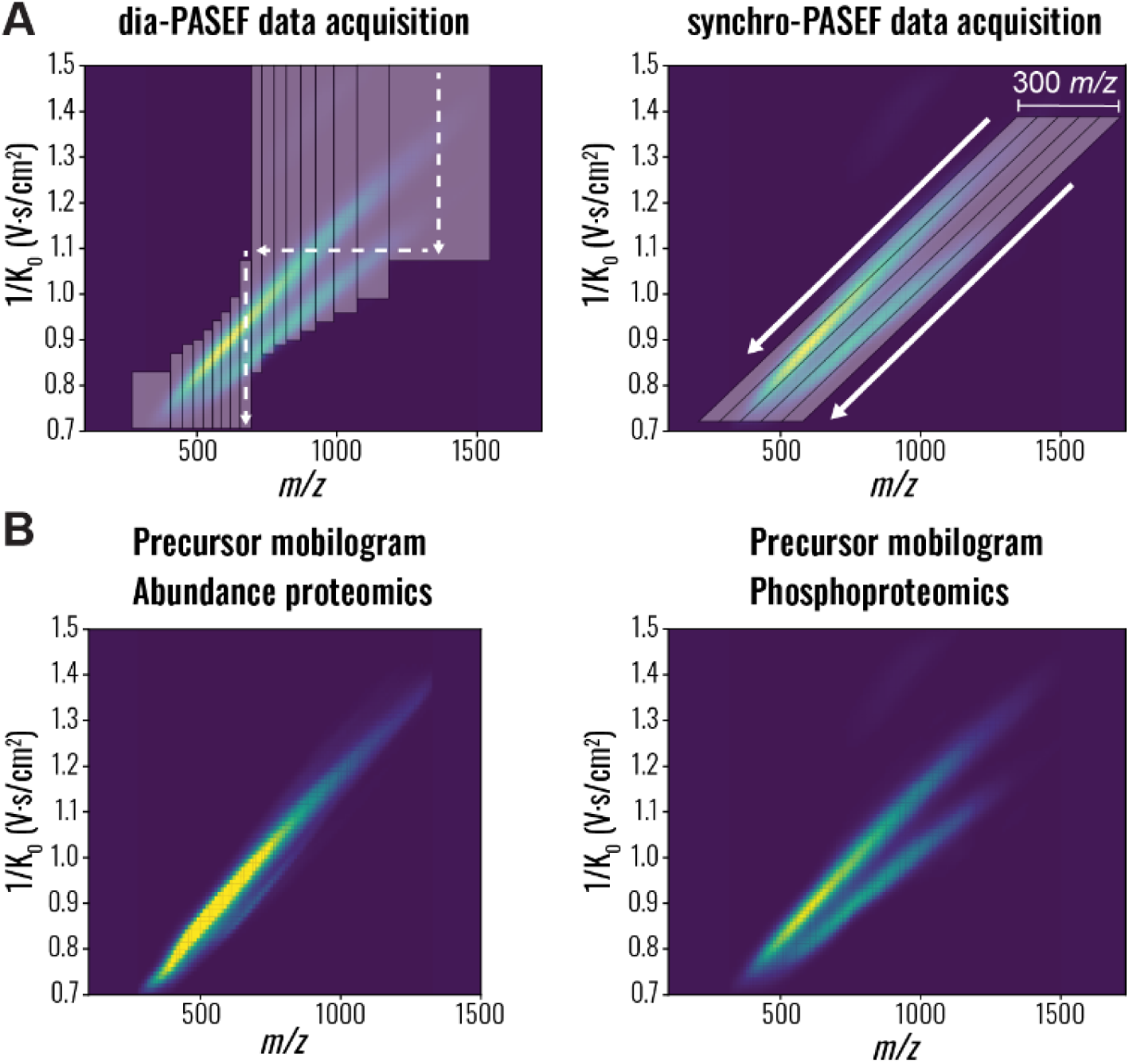
Benchmarking synchro-PASEF method performance for phosphoproteomics across chromatographic separation lengths. **(A)** Illustration of precursor isolation windows using dia-PASEF (left) and synchro-PASEF (right) across a representative HEK293-derived phosphopeptide mobilogram. Dia-PASEF utilizes sequential stepped isolation windows which sample discrete *m/*z regions while scanning through ion mobility space. Synchro-PASEF utilizes continuously stepped isolation windows which track the phosphopeptide precursor ion cloud. Colored density represents the typical precursor distribution, while shaded regions indicate isolation windows. **(B)** Kernel density estimates of precursor ions identified in representative bulk proteomics samples (left) compared to the broader distribution of phopshopeptides (right) across the mobilogram. These distributions illustrate the motivation behind acquisition parameter optimization for phosphoproteomics.

More recently, diagonal scanning strategies such as synchro-PASEF have been introduced in which the TIMS device and quadrupole are ramped simultaneously, generating diagonal isolation windows that track the diagonal distribution of peptide precursors across mobility and m/z space, leading to improved ion utilization. (19–21) Initial reports suggest that the more efficient scanning of synchro-PASEF improves quantitative performance while maintaining proteome depth, particularly in fast chromatographic workflows and with low sample loading amounts. (22, 23) Fewer diagonal isolation windows are required to cover the precursor ion distribution, resulting in shorter MS cycle times and increased precursor sampling frequencies. Analogous to established ion mobility down-sampling practices, higher precursor sampling frequencies enable down-sampling in the retention time dimension by summation (RTsum), improving signal-to-noise, peptide identification rates, and quantitative reproducibility. (22) However, this work was conducted exclusively on global abundance proteomics samples, and a systematic evaluation of synchro-PASEF performance benchmarked to established dia-PASEF strategies across different sample types, acquisition parameters, and gradient lengths has not yet been performed. Phosphoproteomics, which is widely used for example to study kinase signaling (24–27) and drug mechanisms of action (28–30), presents a relevant test case as phosphorylation broadens and reshapes the precursor ion cloud into a bimodal distribution (**Fig. 1B**), which might impact isolation window design, analysis depth, and reproducibility by synchro-PASEF. (31)

In this study, we present a comprehensive framework to evaluate synchro-PASEF data acquisition on the timsTOF HT platform for high-throughput phosphoproteomics by benchmarking its performance against dia-PASEF. Using phosphopeptides enriched from cell lysates, we first systematically optimized the total isolation width and number of diagonal ramps across three sub-hour gradient lengths around the bimodal phosphopeptide precursor ion cloud. We then demonstrated that retention time summation (RTsum) broadly improves both identification depth and quantitative reproducibility across all synchro-PASEF and dia-PASEF methods tested. Finally, we applied our optimized synchro-PASEF and dia-PASEF method to model phosphopeptide dose-response relationships downstream of β2-adrenergic receptor (B2AR) activation. Previous phosphoproteomic dose-response studies have relied primarily on data- dependent acquisition (DDA) coupled with tandem mass tag (TMT) labeling (28, 29) or an orthogonal gas-phase ion mobility filtering device (32) which require labeling reagents and constrain the number of conditions that can be directly compared within a single multiplexed set. We demonstrate that both synchro-PASEF and dia-PASEF capture concentration-dependent phosphorylation dynamics and recover known protein kinase A (PKA)-dependent signaling events downstream of B2AR activation establishing label-free DIA phosphoproteomics as a flexible and scalable alternative for quantitative dose-response studies. Together, this work provides practical guidelines for synchro-PASEF method design in phosphoproteomics, demonstrates that synchro-PASEF and dia-PASEF yield comparable biological insights in a well-characterized signaling model, and establishes retention time summation as a broadly beneficial strategy for PASEF-based workflows.

## Experimental Procedures

### Experimental Design

#### HEK293 Cell Culture

HEK293 cells were obtained from the American Type Culture Collection (ATCC, CRL-1583), cultured in Dulbecco’s modified Eagle’s medium (DMEM, Corning), supplemented with 10% fetal bovine serum (Thermo Fisher) and 1% penicillin/streptomycin (Corning), and maintained at 37°C in a 5% CO2 humidified incubator.

#### Isoproterenol Dose-Response Treatment and Cell Lysis

HEK293 cells were seeded in 12-well plates coated with poly-D-lysine 24 hours before isoproterenol treatment. Cells were seeded at a density of 3.5×10^5^ per well, aiming at a cell confluency of around 80% for the isoproterenol treatment. Isoproterenol stock solutions were generated at 5x concentration by serial dilutions (**Table 1**) and then added to the cell at a 1:5 dilution to reach the final concentrations.

Following 1 min isoproterenol treatment, cells were rapidly washed three times with ice-cold Dulbecco’s phosphate-buffered saline (PBS, Corning) and lysed directly in the plate using 250 uL 4% Sodium desoxycholate (Sigma) in 100 mM Tris pH8, collected in a tube, and the lysates were incubated at 95 degrees C for 10 min. The lysates were cooled to room temperature and sonicated by microtip probe sonication (Thermo Fisher). Sonication parameters were set to a total runtime of 10 seconds with pulses of 1 second on and 1 second off at an amplitude of 40%, each sample was sonicated twice. Following sonication, protein concentration was determined using the Pierce™ Rapid Gold BCA Protein Assay Kit (Thermo Fisher Scientific).

#### Protein Aggregation Capture and Protein Digestion

For each sample, 150 µg of total protein lysate was used for further processing, all samples were normalized to 150 µg total protein in 200 µl lysis buffer. Reduction and alkylation was performed using tris-(2-carboxyethyl) (TCEP) (10 mM final) and 2-chloroacetamide (CAA) (40 mM final) for 30 minutes at 37°C. Protein digestion was performed according to an adapted version of the Protein Aggregation Capture (PAC) on a KingFisher Flex System (Thermo Scientific) with MagReSyn Hydroxyl beads (ReSyn Biosciences) followed by on-bead digestion. (33) KingFisher deep-well plates were prepared for washing steps, containing 1 ml of 100% Acetonitrile (ACN) or 70% Ethanol (EtOH). For each sample, 100 μl of digestion solution (50 mM ammonium bicarbonate (ABC)-buffer) was prepared and transferred to a regular KingFisher plate. For the samples, the 200 µl cell lysates were mixed with 466 µl 100% ACN to achieve a final concentration of 70% ACN. The storage solution from the Hydroxyl beads was replaced with 70% ACN and an equivalent of 30 µl beads at 20 mg/ml (600 µg beads for each sample at a protein to bead ratio of 1:4) were added to the samples. Protein aggregation was carried out in two steps of 1 min mixing at medium speed, followed by a 10 min pause each. Sequential washes were performed in 2.5 min at a slow speed without releasing the beads from the magnet. Trypsin (Promega) and LysC (FUJIFILM Wako Chemicals) were added at a 1:50 (enzyme:protein w:w) ratio and digested for 6 hours at 37°C. Following digestion, 10% trifluoroacetic acid (TFA) was added to each sample to a final pH ∼2. For the protein abundance samples, an equivalent volume to 10 µg per sample was desalted using BioPureSPN PROTO 300 C18 Macro Spin Column (Nest Group). Each column was activated with 200 μL methanol, washed with 3 x 200 μL 80% acetonitrile (ACN)/0.1% TFA, then equilibrated with 3 x 200 μL of 2% ACN/0.1% TFA. Following sample loading, cartridges were washed with 3 x 200 μL of 2% ACN/0.1% TFA, and samples were eluted with 2 x 200 μL 50% ACN/0.1% TFA. Samples were dried by vacuum centrifugation and resuspended in 0.1% formic acid (FA) before mass spec measurements. The remainder of the digested samples (140 μg/sample) was subjected to phosphopeptide enrichment.

#### Phosphoproteomics Sample Preparation

Phosphopeptide enrichment was performed on a KingFisher Flex System (Thermo Scientific) using MagReSyn zirconium-based immobilized metal affinity chromatography (Zr-IMAC HP) beads (ReSyn Biosciences) as described previously. (33) Dried peptides were resuspended in 200 μl loading buffer (80% ACN, 5% TFA, 0.1 M glycolic acid) and transferred to a KingFisher 96 deep-well plate. Additional KingFisher plates were prepared containing 500 μl of loading buffer, 500 μl of washing buffer 2 (60% ACN 1%TFA 200mM NaCl), 500 μl of washing buffer 3 (60% ACN 1% TFA), and 500 μl of washing buffer 4 (500 μL of HPLC-grade water) each. For each sample, 15 μl of beads (20 mg/ml) were washed twice in loading buffer, resuspended in 200 μl loading buffer, and added to a KingFisher plate. For elution, 200 μl of elution buffer (1% NH4OH) was prepared and transferred to KingFisher plates. Beads were washed in the loading buffer for 5 min, incubated with the samples for 20 min with mixing at medium speed, and subsequently washed in loading buffer, washing buffer 2, washing buffer 3, and washing buffer 4 for 2 min with mixing at fast speed. Phosphopeptides were eluted from the beads by mixing with the provided elution buffer for 10 min at a fast speed. Following elution, the samples were acidified with 10% TFA to pH ∼2 and desalted using BioPureSPN PROTO 300 C18 Micro Spin Column (Nest Group) as described above.

#### Liquid Chromatography – Mass Spectrometry

Phosphoproteomic samples were analyzed using a TimsTOF HT mass spectrometer (Bruker) coupled to a Vanquish Neo ultra high-pressure liquid chromatography system (Thermo Fisher Scientific) via a CaptiveSpray2 nanoelectrospray source. Samples were reconstituted in 0.1% formic acid and loaded onto an IonOpticks 15 cm x 75 μm I.D. Aurora CSI column. Mobile phase A consisted of 0.1% FA, and mobile phase B consisted of 0.1% FA/80% ACN. Peptides were separated at a flow rate of 500 nL/min using a nonlinear gradient increasing buffer B from 2% to 80% over 8-, 23-, or 52-minutes followed by a set 7-minute column wash and equilibration. All MS experiments acquired data in either dia-PASEF or VistaScan mode (for synchro-PASEF methods) using a 75 ms TIMS ramp and utilized a TIMS precursor MS window covering an *m/z* range of 100-1700 Da and a 1/K0 range of 0.72-1.50. For all data acquisition methods, this TIMS precursor MS scan preceded all dia-PASEF or synchro-PASEF MS^2^ scans. All dia- and synchro-PASEF methods tested were acquired in biological triplicate. The dose-response dataset was collected in biological quadruplicate.

#### diaPASEF and synchro-PASEF Method Construction

For phosphoproteomics samples, a preliminary dia-PASEF method (denoted as *Static Window dia-PASEF)* was constructed using the TimsControl method editor by drawing a polynomial around the bimodal phosphopeptide precursor distribution. The total number of windows was set to 9, the 1/K0 range was set to 0.72-1.50, window *m/z* isolation widths were set equal, and a 1 Da overlap was used between overlapping window segments. Following data processing using Spectronaut (refer below), the resulting phosphopeptide identifications were then fed into the dia-PASEF and synchro-PASEF method optimization software *py_diAID*. (17, 19) *Py_diAID* enables the facile generation of dia-PASEF methods with variable width isolation windows and synchro-PASEF methods from a set of previously identified peptide precursors. Using the reported identifications from a previously produced human phosphoproteomics dataset, we generated a second benchmark dia-PASEF method (denoted as *Variable Window dia-PASEF)* using 9 tims MS^2^ ramps. We set the *m/z* range from 250-1500 *m/z* and each window to be split once with a 1 Da overlap between overlapping window segments for all dia-PASEF methods. The same precursor collection was used to produce all synchro-PASEF methods; this collection is used to scale and center all diagonal synchro-PASEF windows over the distribution of identified precursors in *m/z*–1/K0 space. The 1/K0 range was fixed from 0.72-1.50. To ensure reproducible coverage of peptide precursors when varying diagonal window count from 2 to 12 windows, the number of diagonal ramps were split evenly over a predefined total *m/z* width (200, 300, or 400 Da total) centered on the peptide ion cloud. Total *m/z* width was slightly adjusted beyond the nominal *m/z* range for each window count to ensure even unit division across all windows for the purposes of preventing rounding errors in generated window schema. No *m/z* overlap between the synchro-PASEF isolation windows were used.

### Statistical Rationale

#### Proteomics Statistical Analysis

All resulting mass spectrometry data were analyzed with Spectronaut (Biognosys, v19.9) using direct DIA analysis for the identification and quantification of the resulting dia-PASEF and synchro-PASEF datasets. (34) The resulting raw data files were binned by data acquisition method and sample type and serially processed using Spectronaut’s command-line interface in batch mode using an in-house C# script. As the Spectronaut data processing algorithm differs between dia-PASEF data and synchro-PASEF data, search parameters were slightly modified depending on the mode of acquisition. The following changes to the default BGS factory settings were made for both dia-PASEF and synchro-PASEF datasets. Cross-run normalization was turned off. Acquired data were searched against the UniProt canonical human proteome (downloaded October, 2024) and the BGS contaminant database. (34, 35) Carbamidomethylation of cysteine (C) was used as a fixed modification, and protein N-terminus acetylation and methionine oxidation were set as variable modifications. For phosphoproteomics experiments, phospho (STY) was also set as a variable modification with a maximum of 2 variable modifications. The PSM, peptide, and protein group false discovery rates were set to 0.01 with no missing value imputation enabled. For synchro-PASEF searches *diaPASEF Pre-Processing* was set to Legacy (Spectronaut 18) and *IM Sampling Reduction* was set to a factor of 3 as suggested by Below, *et al*. (22) For dia-PASEF searches *diaPASEF Pre-Processing* was set to Automatic, and *IM Sampling Reduction* was set to the default factor of 7. Retention time sampling reduction (RTsum factor) was set to 1 unless otherwise indicated in the main text. For dose-response experiments, samples acquired using dia-PASEF and synchro-PASEF methods were batched by acquisition technique, each searched independently, with samples within each batch analyzed together using the Spectronaut GUI and the settings described above.

Following pre-processing the resulting peptide fragment ion intensities were exported from Spectronaut in *tsv* format and were imported into R (v 4.5.1) for further analysis using the MSstatsPTM framework. (36, 37) Default settings were used unless otherwise indicated. After importing transition-level data from Spectronaut into the R environment, peptides from known contaminant proteins were filtered and the dataset was filtered to only retain phosphopeptides which had a 75% localization probability in at least one sample. Following, these data were converted to the standard MSstatsPTM format using the *SpectronauttoMSstatsPTMFormat* function using a qvalue_cutoff of 0.01. These data were summarized to phosphosite-level abundances using the MSstatsPTM dataProcess function (2.4.4). Data were normalized using the *equalizeMedians* argument and missing value imputation was disabled.

The phosphoproteomic dose-response analysis was conducted using the MSstatsResponse package (pre-release 0.99.2). (38) Briefly, class I phosphosite abundances for the selected dia-PASEF and synchro-PASEF methods were imported into the R environment and were ranked by concentration of isoproterenol. Any phosphosite with a response value of 0 was set to *NA*. Each dose-response series was fit to an isotonic regression model using the doseResponseFit function with the appropriate increasing or decreasing directionality constraint. An F-test was performed to assess the significance of the monotonic dose-response curve against the null hypothesis of constant phosphosite abundance across all doses, and resulting p-values underwent FDR correction using the Benjamini-Hochberg procedure. The log2 fold-change of each curve was calculated using the maximum deviation from the control, and thresholds of q ≤ 0.05; log2FC ≥ ±1 were used to determine significantly responding phosphosites across the dose-response series. IC50 and EC50 values were calculated using the predictIC50 function by setting *expected_response* to 0.5 and 1.5 respectively. Kinase enrichment analysis was conducted using the Kinase Library python package which predicts kinase-substrate affinities from positional peptide scanning array assays instead of using prior knowledge. (39, 40) 11-mer amino acid motifs were extracted for each phosphosite from the source FASTA file. Fisher enrichment analysis was then conducted using thresholds of log2-fold change >= ±1, q-value <= 0.05, and a predicted kinase percentile rank threshold of 10.

### Data Availability

The raw MS data, the Spectronaut projects, R notebooks, and the quantitative data tables have been deposited to the ProteomeXchange Consortium via the MassIVE partner repository with the dataset identifier MSVXXXXXXXXX. (41) Quantitative results are provided as exported report files within the repository. A sample-to-raw-file mapping table detailing the correspondence between raw data files, experimental conditions, and figures are included in the repository to facilitate data interpretation. R notebooks can also be accessed via GitHub (https://github.com/HuttenhainLab). The saved search files from Spectronaut can be viewed with the Spectronaut Viewer (www.biognosys.com/spectronaut-viewer).

## Results

### Generation and optimization of synchro-PASEF methods for phosphoproteomic samples at different gradient lengths

We sought to first establish the baseline performance of synchro-PASEF data acquisition methods using a representative set of phosphoproteomic samples and chromatographic separation lengths commonly used in high-throughput proteomic studies (**Fig. 2A, B**). While previous studies optimized their synchro-PASEF methods around the monomodal precursor ion distribution of unmodified tryptic peptides, phosphoproteomic peptides produce a wider bimodal ion cloud distribution that may require different acquisition parameters (**Fig. 1B**). To optimize dia-PASEF and synchro-PASEF fragmentation windows around the wider bimodal ion cloud, we generated a representative set of phosphoproteomics samples from HEK293 cell lysate in biological triplicate (**Fig. 2A**). These samples underwent initial benchmarking using two distinct dia-PASEF methods, one generated using the Bruker timsControl window editor with fixed-width isolation windows (denoted as *Static Window dia-PASEF*) and one optimized using the *py_diAID* software with variable-width isolation windows (denoted as *Variable Window dia-PASEF)* (**Fig. 2C, D**). The resulting phosphopeptide precursor identifications from these dia-PASEF runs were then utilized to generate an array of synchro-PASEF methods via *py_diAID* which varied in the number of fragmentation windows (5, 7, or 9 total) and spanning a total *m/z* width (300, 400, or 500) (**Fig. 2E**). These parameters represent a trade-off between MS cycle time, precursor ion cloud coverage, and spectral complexity from co-isolation of multiple precursors.

**Figure 2.**
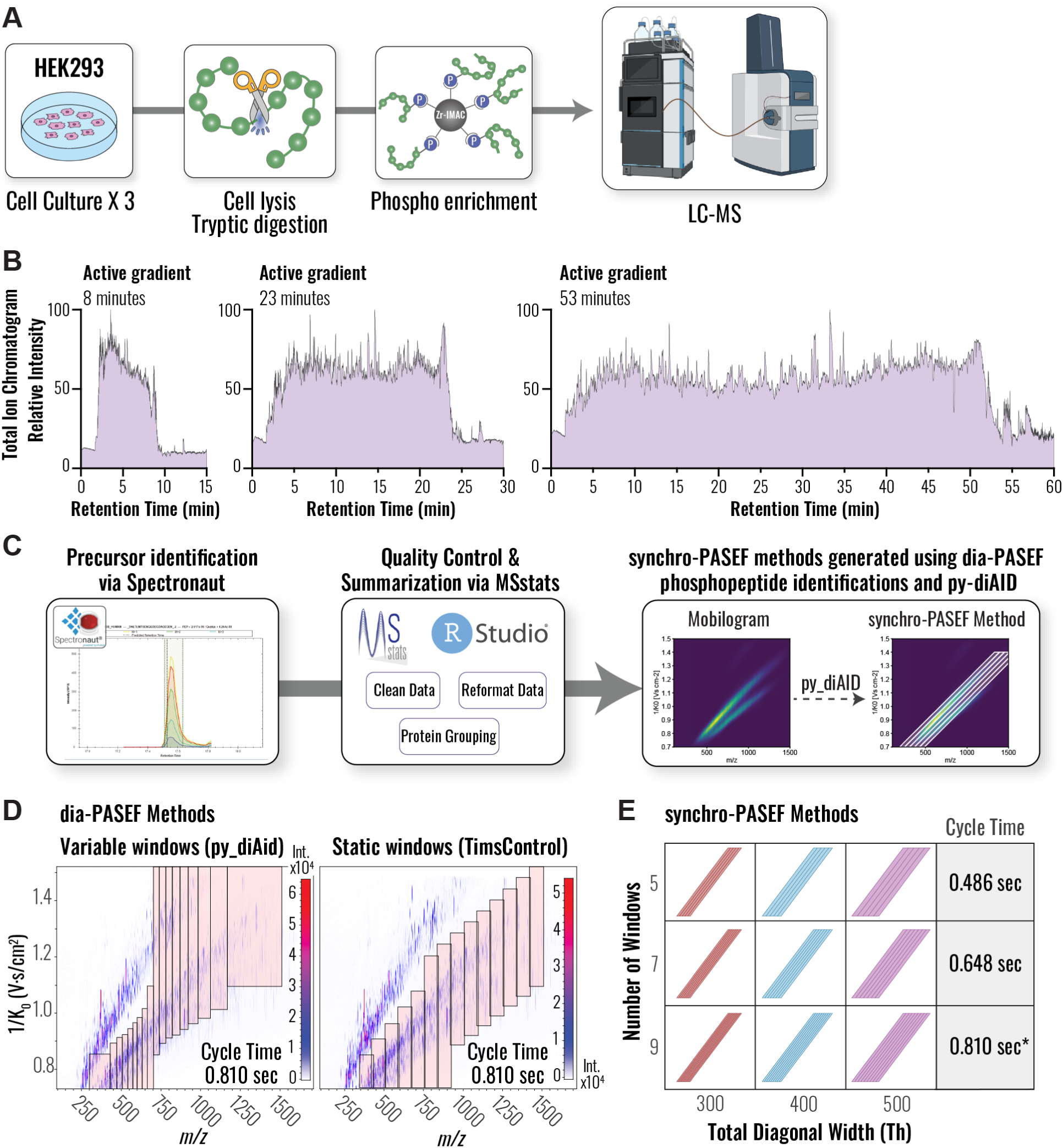
Synchro-PASEF method design guided by phosphopeptide precursor ion cloud distributions. **(A)** Phosphoproteomic samples were derived from HEK293 cell lysate in biological triplicate and were subjected to LC-MS analysis. **(B)** Representative total ion chromatograms for HEK293 phosphopeptides separated using 8-minute, 23-minute, and 53-minute active chromatographic gradients. **(C)** Representative phosphopeptide precursor ion clouds derived from HEK293 cell lysates analyzed in biological triplicate using dia-PASEF acquisition to derive representative phosphopeptide precursor ion clouds. Raw MS data was processed using Spectronaut and MSstatsPTM, and the resulting precursor-level data was used to generate synchro-PASEF methods using the py_diAID software. **(D)** Two dia-PASEF benchmark methods: fixed-width isolation window dia-PASEF (via TimsControl) and variable-width isolation windows dia-PASEF (via py_diAID). **(E)** Array of synchro-PASEF methods systematically varying in total isolation width (300, 400, or 500 m/z) and the number of diagonal windows (5, 7, or 9), with cycle time driven by window count.

Following method generation, we analyzed biological triplicate injections of phosphopeptides from HEK293 lysates using 8-minute, 23-minute, and 53-minute reverse phase separations. Cycle times ranged from 0.49 seconds (5 windows) to 0.81 sec (9 windows) for synchro-PASEF methods. Gradient-dependent data points per peak ranged from average precursor sampling frequencies of 4.3 data points per peak for the 8-minute gradient and 10.6 for the 53-minute gradient. We processed each method’s set of biological replicates separately using Spectronaut and MSstatsPTM to determine the number of identified class I (≥75% localization probability) phosphopeptides (**Fig. 3A**) and phosphosites (**Fig. 3B**) and calculate coefficients of variation at the phosphosite level (**Fig. 3C**).

**Figure 3.**
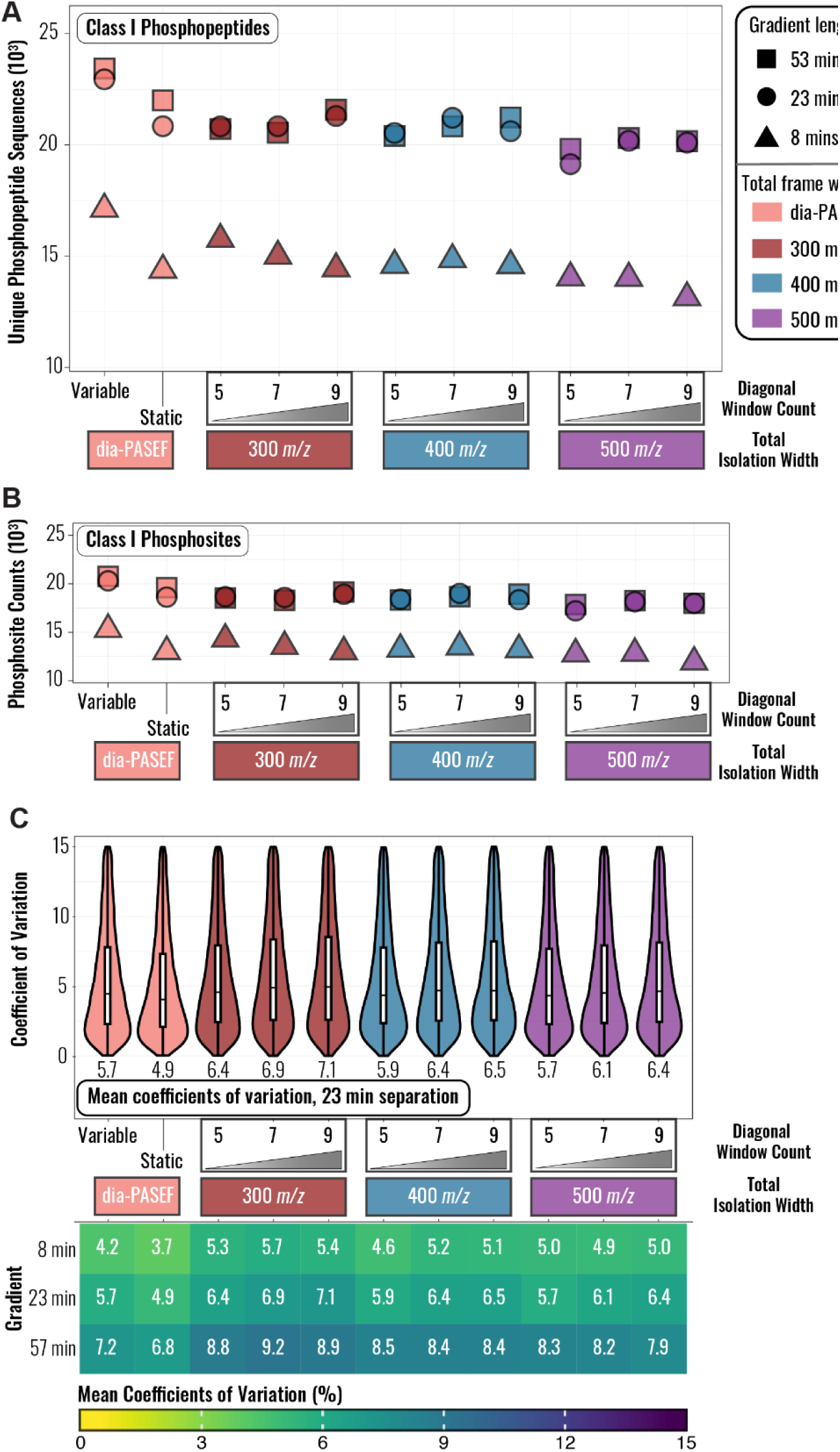
Phosphopeptide identification depth and quantitative reproducibility across dia-PASEF and synchro-PASEF acquisition methods and gradient lengths. **(A)** Class 1 localized phosphopeptide counts across all acquisition methods and gradient lengths. **(B)** Count of class I phosphosites across all acquisition methods and gradient lengths. **(C)** Distribution of phosphosite coefficients of variation for all methods using the 23-minute separation (top). Mean CVs are indicated by labels below the violin plots. Heatmap visualizes mean phosphosite coefficients of variation across all biological replicates for each gradient length and MS acquisition method (bottom).

Synchro-PASEF methods generally produced fewer class I phosphopeptide and phosphosite identifications compared to the variable window dia-PASEF benchmark method regardless of separation length (**Fig. 3A, B**). However, for the 8- and 23-minute gradients, several of the synchro-PASEF methods with 300-400 Da total isolation width outperformed the static window dia-PASEF benchmark (**Fig. 3A, B**). Because approximately 80% of identified phosphosites mapped to a single phosphopeptide, trends at the phosphopeptide and phosphosite levels were largely concordant, and we focus subsequent discussion on phosphosite-level results. (**Fig. 3A, B**). Extending the separation from 23 to 53 minutes provided negligible benefits for most methods, yielding at most minor gains in identifications alongside increased phosphosite-level coefficients of variation (CVs) (**Fig. 3C**).

For the 23- and 53-minute gradients, increasing the number of isolation windows improved identification counts by up to 8% due to improved precursor selectivity. This trend reversed for the 8-minute gradient, where fewer isolation windows - and correspondingly shorter cycle times - were required to adequately sample the compressed elution profiles for reliable identification and quantification (**Fig. 3A, B**). The two best performing synchro-PASEF methods in the 23-minute gradient used 9 or 7 diagonal windows over a total isolation width of 300 and 400 m/z, respectively; both produced ∼21,350 class I phosphopeptides and ∼19,000 phosphosites, slightly exceeding the static window dia-PASEF benchmark. The variable window dia-PASEF benchmark with variable-width windows produced ∼12% more phosphopeptides than the best synchro-PASEF method. In terms of quantitative reproducibility, synchro-PASEF CVs tended to be higher than those for dia-PASEF methods, and CVs generally increased with longer gradients across all methods. Among synchro-PASEF methods, CVs decreased with increasing total isolation width, with no discernable trend in CVs related to the number of diagonal ramps. In summary, the variable window dia-PASEF method produced the highest identification counts, whereas the static window dia-PASEF method achieved the highest quantitative reproducibility, and the best synchro-PASEF configurations matched or slightly exceeded the static window benchmark in identifications.

### Retention-time summation improves phosphopeptide identifications and quantitative reproducibility across all PASEF methods

As synchro-PASEF’s shorter cycle times increase the number of data points sampled per chromatographic peak (**Fig. S1**), we next asked whether this oversampling could be leveraged through retention time summation (RTsum) to improve phosphopeptide identification and quantitative reproducibility. (22) We generated a second phosphoproteomic dataset from HEK293 cell lysates using the 23-minute chromatographic separation, fixing the total isolation width at 400 m/z - based on the best performing synchro-PASEF method in the initial dataset (**Fig. 3**) - while varying the number of diagonal isolation windows from 2-11 ramps (**Fig. 4A**). Both dia-PASEF benchmark methods (fixed and variable window) were included as comparison. Consistent with our initial findings, dia-PASEF methods generally produced higher class I phosphopeptide and phosphosite identification counts than synchro-PASEF methods prior to RTsum (**Fig. 4B**).

**Figure 4.**
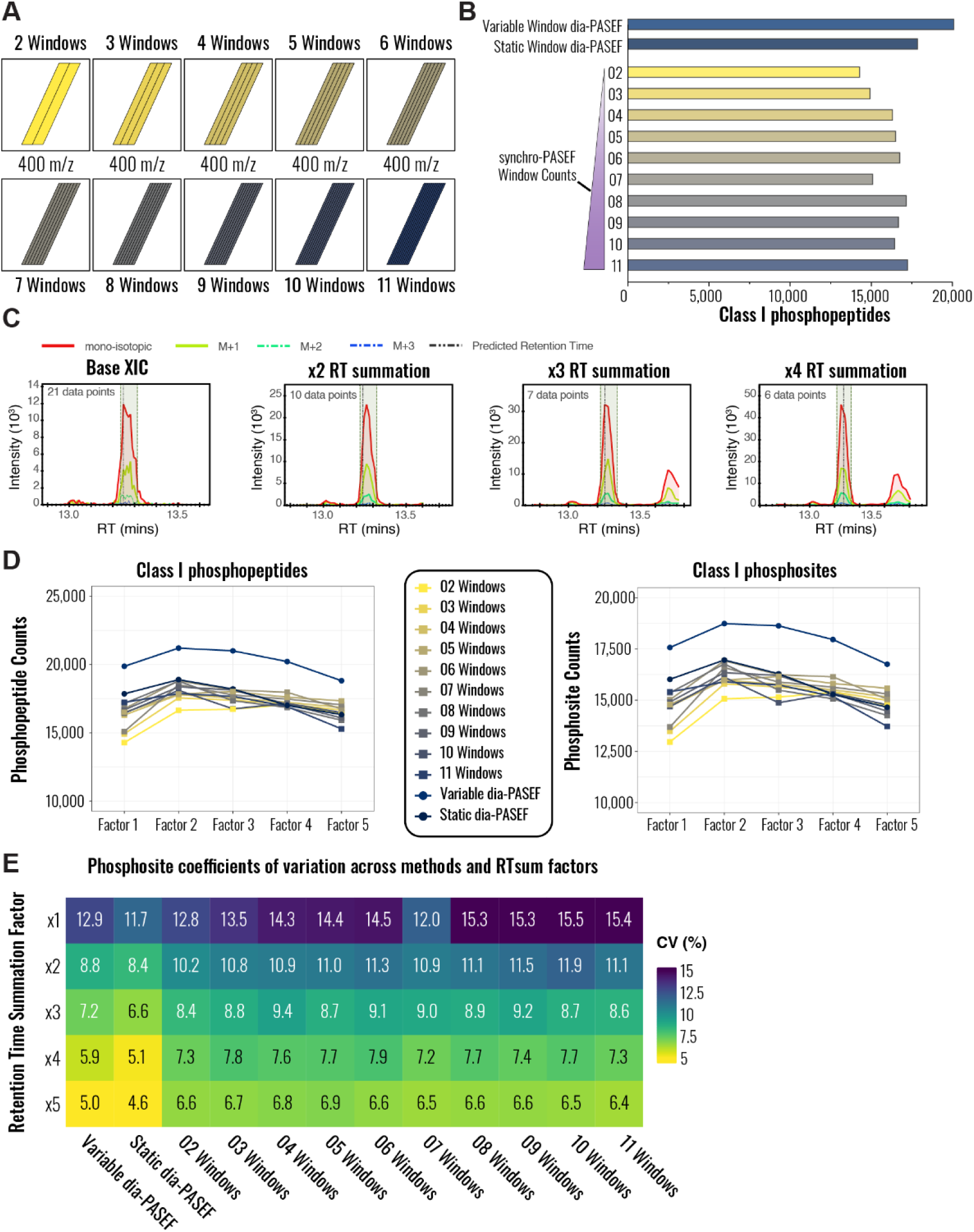
Retention time summation broadly improves phosphosite identification depth and quantification reproducibility across PASEF acquisition methods. **(A)** Synchro-PASEF methods were generated with a fixed total isolation width of 400 *m/z* while systematically varying the number of isolation windows. (**B**) Class I phosphopeptide identification counts produced by each acquisition method without retention time summation. **(C)** Effect of retention time summation exemplified for the extracted ion chromatogram of the phosphopeptide [*p*SIPLSIK]^+2^ acquired with a 2-window synchro-PASEF method. Higher RTsum factors aggregate the signal from multiple scans into a single pseudo-scan. **(D)** Class I phosphopeptide (left) and phosphosite (right) identification counts across all methods and RTsum factors 1-5. **(E)** Heatmap of mean phosphosite coefficients of variation for each combination of acquisition method and RTsum factor.

To investigate how RTsum impacts phosphoproteomic identifications and quantitative reproducibility, we iteratively reprocessed all raw data using RTsum factors starting from 2, 3, 4, and 5, while preserving all other Spectronaut settings. As illustrated for the phosphopeptide [*p*SIPLSIK]^+2^ acquired using a 2-window synchro-PASEF method (**Fig. 4C**), increasing RTsum factors progressively improved signal-to-noise, while proportionally decreasing the number of data points per peak. We observed broad benefits to using RT down-sampling across all acquisition methods (**Fig. 4D, E**). With a RTsum factor 2, every dia-PASEF and synchro-PASEF method demonstrated an increase in class I phosphopeptide and phosphosite identifications–typically 5-8% for most methods, with gains of up to 20-25% for some of the fastest cycling synchro-PASEF configurations (2-3 windows) that provided the greatest degree of oversampling (**Fig. S2**). For these fastest cycling methods, identifications continued to improve modestly at an RTsum factor of 3, but most methods showed diminishing improvements or decreased identification counts with RTsum factors of 3 or higher (**Fig. 4D**). In contrast, quantitative reproducibility improved continuously with increasing RTsum factors: mean phosphosite CVs decreased continuously with increasing RTsum factors, with reductions of up to 50% using an RTsum factor of 5 when relative to no summation (**Fig. 4E**).

These results indicate that applying a small RTsum factor of 2 provides the best trade-off providing both improved identifications and reduced CV. More broadly, applying RTsum during post-processing–scaled appropriately to cycle time–broadly benefits both identification depth and quantitative reproducibility across dia-PASEF and synchro-PASEF phosphoproteomics workflows.

### Synchro-PASEF and dia-PASEF methods yield comparable biological insights downstream of β2-adrenergic receptor activation

Having established acquisition and post-processing parameters for synchro-PASEF phosphoproteomics, we next asked whether both label-free DIA methods, synchro-PASEF and dia-PASEF with RTsum, could model phosphopeptide dose-response relationships, an application previously assessed primarily through TMT-based DDA workflows (references). We chose the β2-adrenergic receptor (B2AR), a well-characterized G protein-coupled receptor (GPCR) whose activation drives cyclic AMP (cAMP)-dependent PKA signaling (42–45), as a model system to evaluate concentration-dependent phosphorylation dynamics across an 11-point isoproterenol dose-response series. HEK293 cells, which endogenously express B2AR, were treated for 1 minute with 11 increasing concentrations of the B2AR agonist isoproterenol (30 pM to 3 µM) in biological quadruplicate (**Fig. 5A**). Following phosphopeptide enrichment, samples were analyzed using a 23-minute separation with either the variable window dia-PASEF benchmark method or a 5 isolation window synchro-PASEF method (400 *m/z* total width), each processed with RTsum factors of 1 and 2. We selected the 5-window synchro-PASEF configuration as this method yielded the one of the highest number of phosphopeptide identifications from the enriched HEK293 lysate before RTsum was enabled. Dose response behavior of the phosphosites were modeled with MSstatsResponse (**Fig. 5B**). (38)

**Figure 5:**
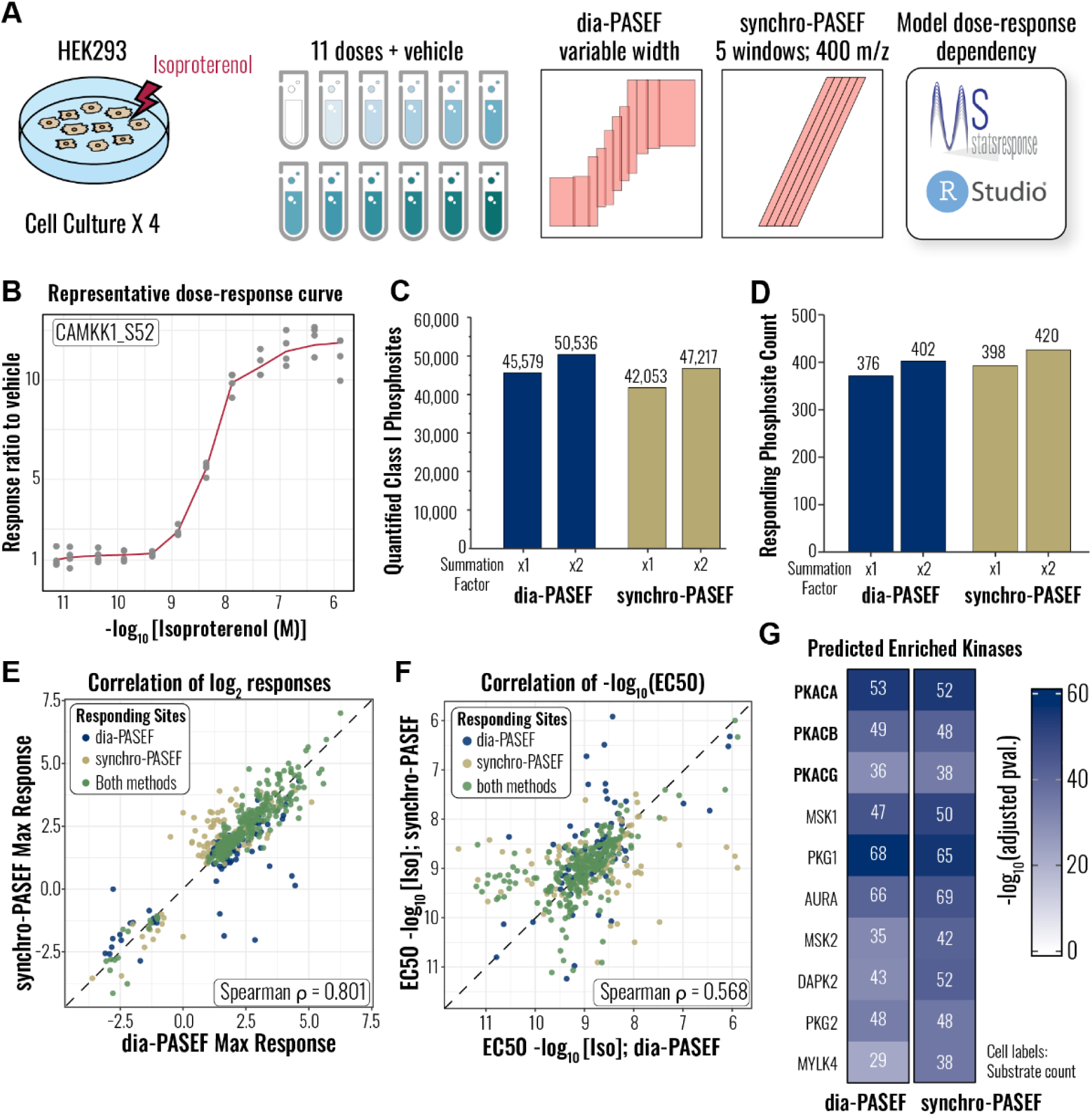
Synchro-PASEF and dia-PASEF methods yield comparable biological insights from β2-adrenergic receptor (B2AR) activation across an isoproterenol dose-response series. **(A)** Overview of the experimental design: HEK293 cells were treated for 1 minute with 11 concentrations of the B2AR agonist isoproterenol (30 pM - 3 µM) in biological quadruplicate. Phosphopeptide-enriched samples were analyzed using either the variable window dia-PASEF benchmark or a 400 m/z-width, 5-window synchro-PASEF method and the dose-response behavior of the resulting data was modeled using MSstatsResponse. (**B**) Representative dose-response curve illustrating phosphosite-level modeling using MSstatsResponse. (**C**) Total counts of quantified class I phosphosites for each acquisition method and RTsum factor. (D) Total counts of significantly responding class I phosphosites for each acquisition method and RTsum factor. (E) Correlation of maximum phosphosite abundance changes (treated vs. vehicle) between dia-PASEF and synchro-PASEF. (F) Correlation of estimated EC50 and IC50 values from modeled dose-response curves between acquisition methods. (G) Kinase-substrate enrichment analysis using the Kinase Library (reference), showing predicted substrate counts for the top 10-scoring kinases across both acquisition methods.

Consistent with earlier results, dia-PASEF produced higher counts of total quantified phosphosites than synchro-PASEF and retention time summation increased this number by 11% and 12% for dia-PASEF and synchro-PASEF, respectively (**Fig. 5C**). However, after phosphosite-level dose-response modeling using MSstatsResponse, synchro-PASEF yielded slightly more significantly responding phosphosites (420) than dia-PASEF (402) (**Fig. 5D, Fig. S3A, B**), with ∼65% overlap between the two methods (**Fig. S3C**). Across the phosphosites found to be significantly responding in either method, we observed good agreement in the maximum phosphosite abundance changes (treated vs. vehicle) between the acquisition methods (Spearman ρ = 0.8, **Fig. 5E**), while the concordance between estimated EC50/IC50 values was weaker (Spearman ρ = 0.57, **Fig. 5F**), likely reflecting the sensitivity of half-maximal response estimation to datapoint density and missing values rather than a systematic bias in either method. A complete heatmap of all responding phosphosites (**Fig. S3D**) shows strong agreement in response trajectories between the two methods, with gaps primarily attributable to phosphopeptides that failed identification or localization in one method.

Non-overlapping phosphosites arose from multiple sources (**Fig. S3C**): approximately 2% of the uniquely responding phosphosites were not sampled because the precursors fell outside the isolation window boundaries of the alternate method; an additional 20% of the phosphosites fell within the sampling regions, but did not pass identification thresholds in Spectronaut (**Fig. S4A**); and the remaining 78% uniquely responding phosphosites were filtered by MSstatsPTM due to low quality or missingness, or failed to reach significance thresholds in MSstatsReponse (**Fig. S4B**).

Finally, we assessed whether the two methods yielded equivalent biological information through kinase enrichment analysis. We performed kinase enrichment analysis using the Kinase Library, which predicts kinase-substrate relationships from *in vitro* positional scanning peptide arrays. (30) We extracted 11-mer motifs for all significantly responding phosphosites and applied Fisher’s exact test using predicted kinase-substrate relationships from this library. Both methods yielded nearly identical counts of predicted PKA substrates (PKACA, PKACB, and PKACG) (**Fig. 5G, Fig. S3E, F**), confirming that synchro-PASEF does not alter the inferred kinase activity landscape despite differences in raw identification counts. Together, these results demonstrate that synchro-PASEF recapitulates the same core biological conclusions as dia-PASEF, supporting its use as a robust alternative for phosphoproteomic studies.

## Discussion

High throughput phosphoproteomics increasingly depends on acquisition strategies that balance proteome depth, quantitative precision, and throughput. Here, we systematically evaluated synchro-PASEF for phosphoproteomic analyses across multiple chromatographic gradient lengths and acquisition configurations, establishing a framework to guide synchro-PASEF method design for phosphopeptide-enriched samples. Although synchro-PASEF’s diagonal scanning strategy and improved ion utilization efficiency is conceptually attractive, our results indicate that its phosphopeptide identification and quantification performance is broadly comparable to - rather than clearly superior to - dia-PASEF methods under the experimental conditions tested. Across all gradients examined, synchro-PASEF methods generally produced fewer localized phosphopeptide and phosphosite identifications than the variable-width dia-PASEF benchmark, though several configurations exceeded the performance of the static-window dia-PASEF method. One explanation for this finding is that variable-width dia-PASEF methods have partially addressed the ion utilization challenge that synchro-PASEF was designed to solve : by matching isolation window specificity to expected phosphopeptide precursor density across the mobilogram, variable-width window methods generated by *py_diAID* already boost precursor coverage efficiency to a degree which attenuates the marginal benefit of diagonal scanning. Additionally, the wider effective isolation width inherent to diagonal windows increases the frequency of precursor co-isolation. This may disproportionately impact the identification of phosphopeptides relative to unmodified peptides. Despite these identification differences, our findings underscore that the benefits of diagonal scanning depend strongly on the balance between cycle time, precursor selectivity, and spectral complexity. Notably, the optimal number of isolation windows varied with chromatographic gradient length: longer gradients benefitted from increased precursor selectivity afforded by more windows, whereas shorter gradients favored fewer windows to preserve adequate sampling frequency for rapidly eluting peaks.

A central finding of this work is that post-acquisition signal processing strategies can significantly influence phosphoproteomic performance independent of acquisition method employed. By exploiting the higher precursor sampling frequencies enabled by shorter cycle times, we demonstrate that retention time summation (RTsum) improves both phosphopeptide identification rates and quantitative reproducibility across all PASEF methods examined. An RTsum factor of 2 consistently increased phosphosite identifications and quantitative precision across all acquisition methods–in some cases by as much as 20–25%–while simultaneously improving quantitative precision. Higher summation factors continued to reduce coefficients of variation even as identification gains plateaued or declined, indicating that the optimal RTsum factor depends on whether the primary analytical objective is identification depth or quantitative reproducibility. These findings are consistent with prior reports that controlled signal averaging in the RT dimensions enhances signal-to-noise by smoothing stochastic fluctuations across neighboring scans. (22) Importantly, the benefits of RTsum were observed for both dia-PASEF and synchro-PASEF datasets, suggesting that RT summation should be broadly incorporated into downstream processing pipelines for PASEF-based phosphoproteomics–particularly in high-throughput workflows where rapid separations increase spectral noise and peak sparsity.

We next assessed whether differences between acquisition strategies meaningfully affect biological interpretation. Using activation of the B2AR and PKA-dependent signaling as a well-characterized model system, we found that synchro-PASEF and dia-PASEF produced highly concordant phosphoproteomic responses across an isoproterenol dose-response series in HEK293 cells. Both methods quantified comparable numbers of significantly responding phosphosites with substantial overlap, similar effect sizes, and nearly identical numbers of predicted PKA substrate counts by kinase enrichment analysis. The small subset of discordant sites arose from stochastic identification differences and downstream statistical filtering rather than systematic divergence in biological signal detection. While maximum response amplitudes were highly concordant between methods (Spearman ρ = 0.8), the weaker agreement in EC50/IC50 estimates (ρ = 0.57) suggests that half-maximal response parameters are more sensitive to missing values and noise at the estimated response intervals. Nevertheless, the strong concordance in responding phosphosite identity, effect size, and kinase activity predictions demonstrates that synchro-PASEF captures signaling dynamics with fidelity comparable to established dia-PASEF workflows. Beyond the method comparison, these results demonstrate that label-free DIA phosphoproteomics can model quantitative dose-response curves for individual phosphosites. Prior phosphoproteomic dose-response studies have predominantly employed label-free or TMT-based DDA approaches (28, 29, 32), which require costly labeling reagents and limit the number of directly comparable conditions to the TMT multiplexing capacity. DIA-based workflows overcome both constraints, enabling flexible experimental designs and direct quantitative comparison across all conditions without batch effects inherent to multi-plex TMT experiments. The concordance between synchro-PASEF and dia-PASEF in recovering PKA-dependent signaling events suggests that either PASEF-based DIA strategy can serve as a scalable, label-free platform for dose-response phosphoproteomics.

For laboratories implementing high-throughput phosphoproteomics on timsTOF instruments, our results provide the following practical guidance: a variable-width dia-PASEF method processed with an RTsum factor of 2 currently offers the best combination of identification depth and quantitative reproducibility for standard-input phosphoproteomic samples analyzed with short chromatographic gradients. Synchro-PASEF with RTsum represents a competitive alternative when flexibility in cycle time tuning is prioritized and may offer distinct advantages where the shortest possible cycle times are required for very brief chromatographic separations.

Several limitations of this study should be noted. First, our benchmarking was performed using input amounts typical of bulk phosphoproteomic workflows. Prior work has suggested that synchro-PASEF can provide advantages at lower sample inputs by improving ion utilization and sensitivity. (22) Under such conditions, differences between acquisition strategies may become more pronounced, and our results may underestimate synchro-PASEF’s potential in low-input or sample-limited settings - including emerging applications such as single-cell phosphoproteomics or limited clinical specimens. Second, our analysis relied exclusively on Spectronaut for data processing; alternative search engines with different scoring algorithms may alter the relative performance of these acquisition strategies. While we focused on synchro-PASEF specifically, the principles established here, particularly regarding the trade-offs between diagonal window design and the benefits of RTsum, are likely generalizable to other diagonal scanning implementations for PASEF-based instruments.

Future work refining acquisition parameters–particularly adaptive window placement guided by the phosphopeptide ion cloud distribution–and integrating RTsum into automated processing workflows will be important to fully realize the potential of synchro-PASEF for large-scale phosphoproteomic studies. Further, systematic evaluation of synchro-PASEF at low sample inputs, including single-cell phosphoproteomics, will be critical to determine whether the theoretical ion utilization advantages translate to practical sensitivity gains under sample-limited conditions where dia-PASEF’s identification advantage may narrow or reverse.

## Acknowledgments

This work was in part supported by the National Institute on Drug Abuse (NIDA) (R01DA056354 to R.H.) of the National Institutes of Health, the Stanford Bio-X Interdisciplinary Initiatives Seed Grants Program (IIP12-21 to R.H.), and the Technology Development Seed Grant Award from the Stanford Beckman Center for Molecular and Genetic Medicine (350788 to R.H.). M.L.G. was supported by NIGMS award T32GM154663.

## Abbreviations

ACN: Acetonitrile
B2AR: Beta-2 adrenergic receptor
BGS: Biognosys
CV: Coefficient of variation
DIA: Data-independent acquisition
EC50: Half maximal effective concentration
FA: Formic acid
FDR: False discovery rate
IC50: Half maximal inhibitory concentration
IMAC: Immobilized metal affinity chromatography
LCMS: Liquid chromatography mass spectrometry
m/z: Mass-to-charge ratio
PASEF: Parallel accumulation-serial fragmentation
PSM: Peptide spectral match
PTM: Post translational modification
RTsum: Retention time summation
TFA: Trifluoroacetic acid
TIMS: Trapped ion mobility spectrometry
TOF: Time of flight

## Notes

### Competing Interest Statement

The authors have declared no competing interest.

