## Supplemental Figures 1-4; Table 1 for "Systematic optimization and benchmarking of synchro-PASEF for high-throughput phosphoproteome profiling"

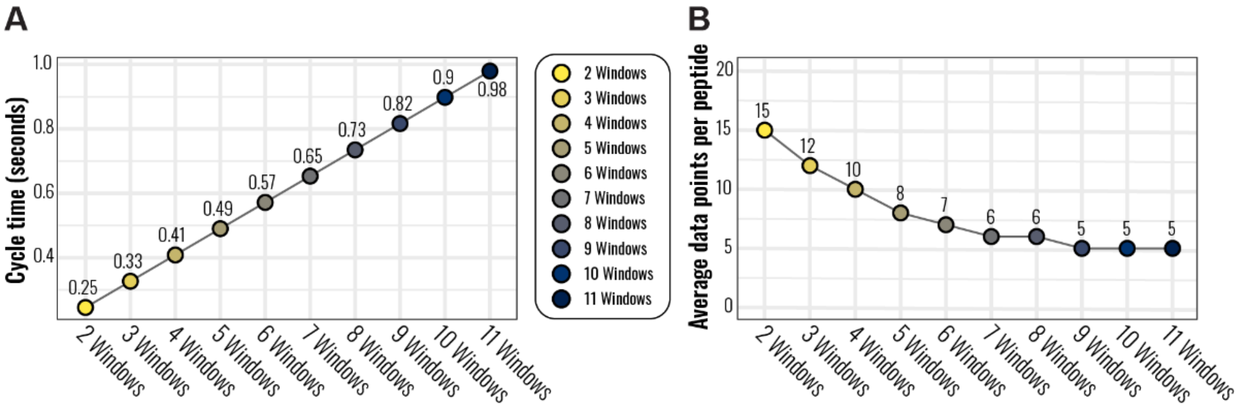


**Supplementary Figure S1 (Related to Fig. 4): Estimated cycle times (A) and mean data points per peak (B) for 23 min separation by number of TIMS ramps.**

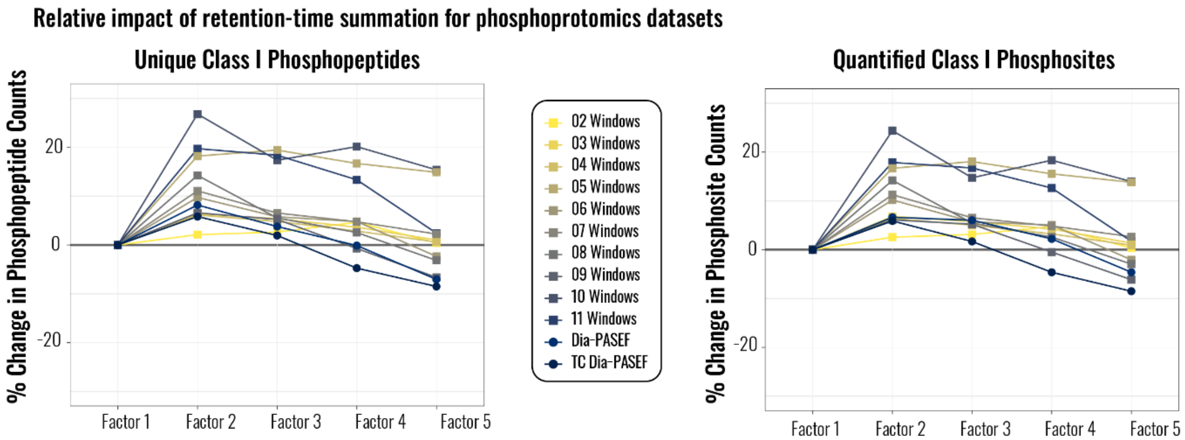


**Supplementary Figure S2 (Related to Fig. 4): Relative impact of retention time summation for phosphoproteomics datasets.** Proportional changes in phosphopeptide (**A**) and phosphosite (**B**) identifications when using retention time summation for phosphoproteomics applications.

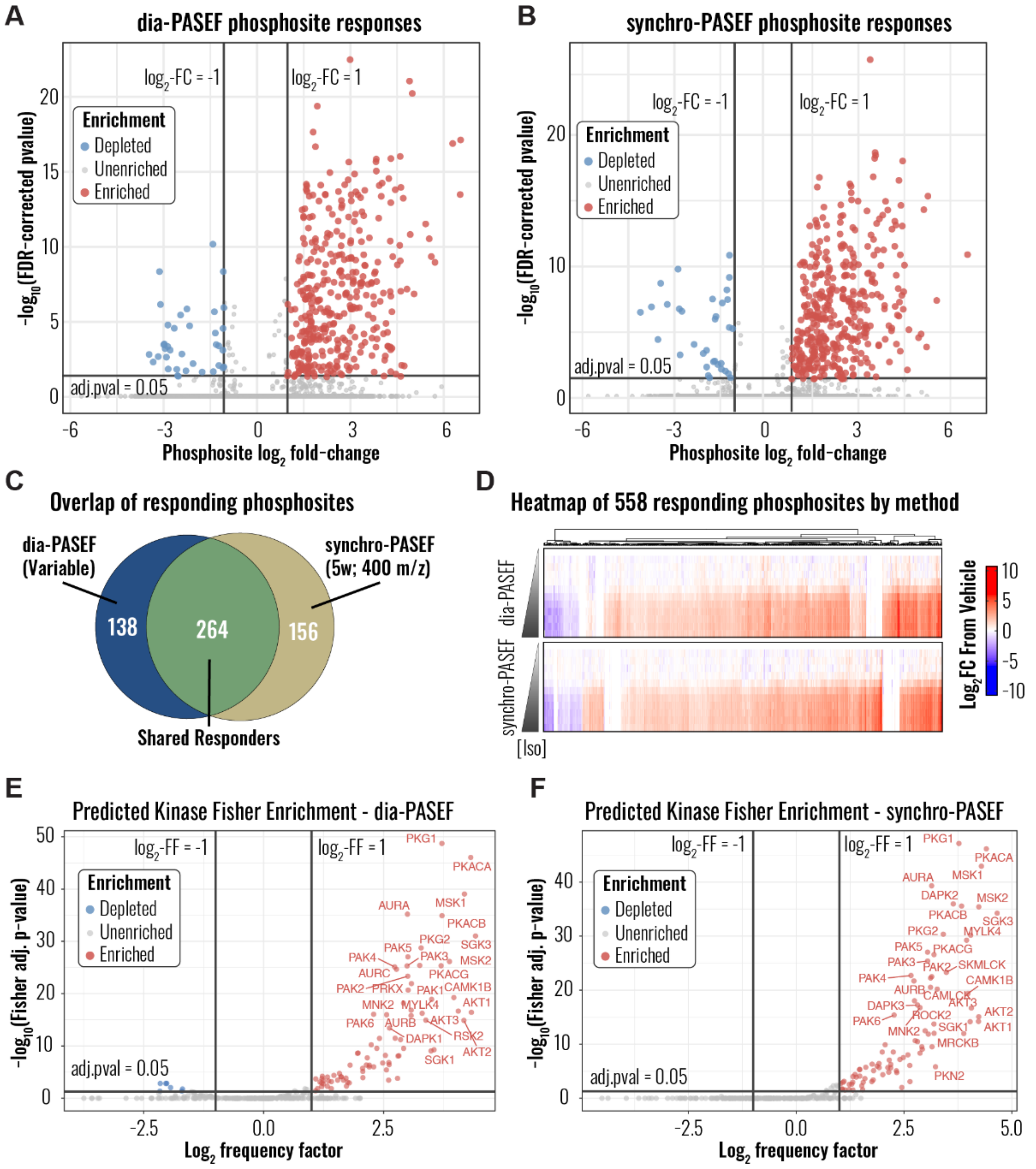


**Supplementary Figure S3 (Related to Fig. 5): Full response profiles for all localized phosphosites from the isoproterenol dose-response series.** Volcano plots of maximal responses and significance of phosphosite dose-response curves for the **(A)** variable-width dia-PASEF and **(B)** 5-window; 400 *m/z* isolation width synchro-PASEF methods. Significance thresholds were set to q ≤ 0.05; $\log_2$FC ≥ ±1. **(C)** Overlap of significantly responding phosphosites between the two acquisition methods. **(D)** Heatmap of all phosphosites significantly responding in at least one acquisition method. **(E)** Kinase-substrate relationship prediction followed by Fisher enrichment analysis for the dia-PASEF dataset and **(F)** the synchro-PASEF dataset.

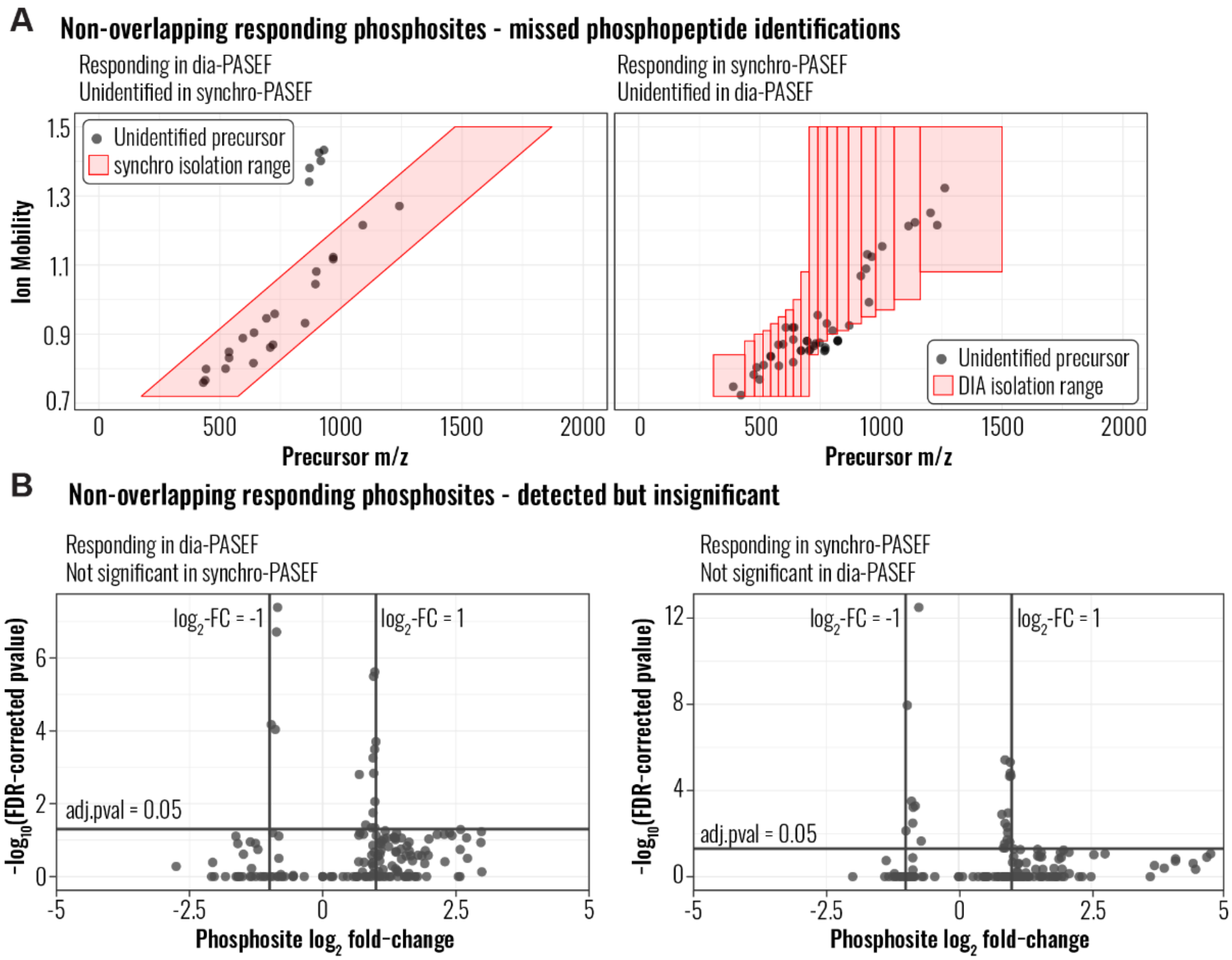


**Supplementary Figure S4 (Related to Fig. 5): Investigation into phosphosites with method-dependent responses.** Phosphopeptides found to be uniquely responding in dia-PASEF or synchro-PASEF methods were cross-mapped to the alternate methods' Spectronaut exports. (**A**) Phosphopeptide precursors from phosphosites uniquely responding in the (left) dia-PASEF method but were not identified in the synchro-PASEF method, and (right) the synchro-PASEF method which were not identified in the dia-PASEF method (or conversely). Isolation windows are overlaid upon the *m/z*-1/Ko values of the missing precursors. (**B**) Phosphopeptides found to be uniquely responding in (left) dia-PASEF or (right) synchro-PASEF methods that were modeled by MSstatsResponse but did not pass significance thresholds.

**Table 1: Isoproterenol dose-response series agonist concentration**

| *Dilution* | *Stock concentration* | *Final agonist concentration* |
|---|---|---|
| Vehicle (No agonist) | 0 nM | 0 nM |
| Isoproterenol stock | 10 mM | N/A |
| Dilution 1 | 15 μM | 3 μM |
| Dilution 2 | 5.0 μM | 1 μM |
| Dilution 3 | 1.5 μM | 300 nM |
| Dilution 4 | 500 nM | 100 nM |
| Dilution 5 | 150 nM | 30 nM |
| Dilution 6 | 50 nM | 10 nM |
| Dilution 7 | 15 nM | 3 nM |
| Dilution 8 | 5 nM | 1 nM |
| Dilution 9 | 1.5 nM | 300 pM |
| Dilution 10 | 500 pM | 100 pM |
